# Heterologous expression, purification and structural features of native *Dictyostelium discoideum* dye-decolorizing peroxidase bound to a natively incorporated heme

**DOI:** 10.1101/2023.05.08.539583

**Authors:** Özlem Kalkan, Sravya Kantamneni, Lea Brings, Huijong Han, Adrian P. Mancuso, Faisal H.M. Koua

## Abstract

The *Dictyostelium discoideum* dye-decolorizing peroxidase (*Dd*DyP) is a newly discovered peroxidase, which belongs to a unique class of heme peroxidase family that lacks homology to the known members of plant peroxidase superfamily. *Dd*DyP catalyzes the H_2_O_2_-dependent oxidation of a wide-spectrum of substrates ranging from polycyclic dyes to lignin biomass, holding promise for potential industrial and biotechnological applications. To study the molecular mechanism of *Dd*DyP, highly pure and functional protein with a natively incorporated heme is required, however, obtaining a functional DyP-type peroxidase with a natively bound heme is challenging and often requires addition of expensive biosynthesis precursors. Alternatively, a heme *in vitro* reconstitution approach followed by a chromatographic purification step to remove the excess heme is often used. Here, we show that expressing the *Dd*DyP peroxidase in 2×YT enriched medium at low temperature (20 °C), without adding heme supplement or biosynthetic precursors, allows for a correct native incorporation of heme into the apo-protein, giving rise to a stable protein with a strong Soret peak at 402 nm. Further, we crystallized and determined the native structure of *Dd*DyP at a resolution of 1.95 Å, which verifies the correct heme binding and its geometry. The structural analysis also reveals a binding of two water molecules at the distal site of heme plane bridging the catalytic residues (Arg239 and Asp149) of the GXXDG motif to the heme-Fe(III) via hydrogen bonds. Our results provide new insights into the geometry of native *Dd*DyP active site and its implication on DyP catalysis.

## 1 Introduction

The *Dictyostelium discoideum* dye-decolorizing peroxidase (*Dd*DyP) is a newly discovered heme peroxidase (Sugano et al., 2007). *Dd*DyP belongs to a new class of DyP-type peroxidases (EC 1.11.1.19), which is different from any other known peroxidases (Kim et al., 1999; Sugano et al., 1999; Sugano et al., 2007; Shrestha et al., 2016). This unique peroxidase family has been shown to perform both H_2_O_2_-dependent oxidation and hydrolytic functions against a wide-spectrum of substrates, ranging from polycyclic dyes, phenolic compounds, sulfides, carotenoids and interestingly lignin biomass, making it a potential candidate for industrial and biotechnological applications including its possible application as bioenergy catalysts as well as biosurfactants in the biodegradation and biotransformation of emerging environmental contaminants (Gan et al., 2022; Rai et al., 2021; Salvachua et al., 2013; Sugano et al., 2021; Xu et al., 2021). This broad substrate specificity is attributed to their unique sequence identity and structural properties (Kim et al., 1999; Yoshida and Sugano, 2023). *Dd*DyP, as other peroxidases, has been found to function in a wide range of pH milieu displaying higher activity at acidic pH with optimal turnover at pH 4.0 and temperatures ranging from 20 to 40 °C (Rai et al., 2021; Xu et al., 2021; Colpa et al., 2014). It shows optimal activity at pH 3.0 against the known DyP-type peroxidase substrate—the anthraquinone-based dye RB4 (Rai et al., 2021).

DyP-type peroxidases share a typical catalytic mechanism with other peroxidases, in which they depend on the H_2_O_2_ in their oxidative catalytic function as illustrated in scheme 1 (Scocozza et al., 2021; Sugano and Yoshida, 2021). The resting state of the enzyme proceeds into compound I intermediate state upon interaction of the heme-Fe(III) with hydrogen peroxide (H_2_O_2_), an oxidizing substrate, forming an oxoferryl porphyrin *π*-cationic radical complex [Fe(IV)═O Por^•^]^+^—a porphyrinoid based radical (Colpa et al., 2014; Scocozza et al., 2021). The release of an electron from compound I leads to the formation of compound II [Fe(IV)═O]^+^ intermediate upon reaction with a reducing substrate giving rise to a radical product, in turn compound II relaxes into the resting state when it reacts with more substrates (Chen et al., 2015; Shrestha et al., 2016; Sugano and Yoshida, 2021). The radical product can then be transformed into various sub-products through a non-enzymatic radical coupling. The redox potential of the DyP-type peroxidases, ranging from -50 mV to +250 mV, and that of the substrate determines the feasibility of enzyme catalysis. Generally, a typical DyP-peroxidase catalysis involves several redox couplings, namely Fe^3+^/Fe^2+^, compound I/Fe^3+^, compound II/compound I and compound II/Fe^3+^ (Shrestha et al., 2016). DyP peroxidases also mediate the hydrolysis of substrates such as anthraquinone dyes, implying that the DyP-type peroxidases are bifunctional enzymes (Colpa et al., 2014). However, the exact mechanism for DyP-type peroxidases and how they perform oxidation and hydrolysis for such a wide range of substrates of different chemical properties remain unclear (Xu et al., 2021; Sugano and Yoshida, 2021).

Despite the importance of the DyP-type peroxidases as mentioned above, their heterologous expression in *Escherichia coli* (*E. coli*) and other expression systems remain challenging. It hampers, for instance, on the large-scale mechanistic investigation owing to the difficulties associated with the biosynthesis and availability of heme *b*, thereby limiting its native incorporation into the apo-proteins (Fiege et al., 2018; Park and Kim, 2021). It was previously shown that *in vitro* reconstitution is needed for obtaining functional *Dd*DyP with the correct heme stoichiometry (Rai et al., 2021). This is an inherently time-consuming process and may result in excess heme and unspecific binding or altering the protein function, making it limited to robust proteins only (Vogel et al., 1999; Denninger et al., 2000; Lemon et al., 2021). Alternatively, heme and iron supplements or heme biosynthetic precursors such as *δ*-aminolevulinic acid (*δ*-ALA) can be used during expression, however this is a highly expensive approach as large amounts of such supplements are needed (Fiege et al., 2018).

Here, we report on the use of *E. coli* OverExpress C43(DE3) strain for the expression and production of the *Dd*DyP peroxidase without heme supplement nor its precursor *δ*-ALA. Furthermore, using X-ray structural analyses, we describe the first crystal structure of native *Dd*DyP peroxidase bound to a natively incorporated heme and demonstrate that the geometry of the heme binding pocket resembles in much detail that of a previously reported cyanide native *Dd*DyP structure, which was prepared following *in vitro* heme reconstitution approach.

## 2 Methods

### 2.1 Overexpression and purification of *Dd*DyP with natively incorporated heme

The gene sequence encoding dye-decolorizing peroxidase from the slime mold *D. discoideum* AX4 (GenBank: EAL70759.1) was codon optimized for *Escherichia coli*, synthesized and subcloned into the *Bam*HI/*Xho*I cloning site in a pGEX-6P1 vector harboring a Human Rhinovirus 3C excision site and a glutathione transferase (GST) tag at the N-terminal region (BioCat GmbH, Germany). The pGEX-6P1-*Dd*DyP construct was transformed into an OverExpress *E. coli* C43(DE3), a chemically competent strain (Sigma-Aldrich, Germany). For purification, a single colony from a freshly streaked plate was inoculated into a Luria-Bertani (LB) or 2×YT enriched media containing 100 μg ml^−1^ final concentration of ampicillin and incubated at 37 °C for overnight (15-16 h). A starter culture was used to inoculate 6× 1L of LB or 2×YT containing 100 μg ml^−1^ ampicillin and incubated at 37 °C or lower temperatures until the optical density (OD_600_) reaches 1.0-1.25 before inducing the expression of *Dd*DyP with 1.0 mM of isopropyl β-D-1-thiogalactopyranoside. The culture was then incubated for additional 7 h or 20 h for expression at 37 and 20 °C, respectively. The cells were harvested by centrifugation with 13,881 ×g for 30 min on an F9-6-×1000 LEX rotor (Thermo Fischer Scientific, Germany) at 4 °C and pellets were stored at -80 °C until used.

For protein purification, frozen cells were thawed using warm tab water (∼40 °C) and diluted with 3-5× of lysis buffer containing 0.05 M Tris-HCl, pH 8.0 and 0.15 M NaCl supplemented with 1.0 mM final concentration of phenylmethylsulfonyl fluoride protease inhibitor or a tablet of EDTA-free protease inhibitor cocktail (Sigma-Aldrich, Germany). The cells were lysed with 35 cycles of sonication at 4 °C on ice using 50% amplitude and 25 sec sonication pulse with 1.5 min interval. Lysate was clarified with centrifugation at 52,400 ×g at 4 °C for 45 min and the supernatant was filtered with a 0.45 μm syringe filter and mixed with 5-10 ml glutathione sepharose high-performance resin pre-equilibrated with lysis buffer, followed by incubation at 4 °C with gentle rotation for 3 h. The mixture was loaded into an empty gravity column and the GST-tagged *Dd*DyP was eluted with 5× column volume of an elution buffer containing 0.05 M Tris-HCl, pH 8.0, 0.15 M NaCl and 15-20 mM L-Glutathione (reduced form). The GST tag was then removed using HRV 3C protease with 1:20 enzyme to protein ratio at 4 °C for overnight followed by passing the mixture through a glutathione sepharose column pre-equilibrated with lysis buffer. Purified protein was characterized with SDS-PAGE and UV-visible spectrophotometry. For crystallization the protein was further purified with gel-filtration using Superdex 75 10/300 Increase column (Cytiva, Sweden). Purified protein was concentrated to 20-30 mg ml^-1^ in lysis buffer and stored at -80 °C until further use. Overexpression, purification and crystallization were carried out the XBI BioLab of the European XFEL facility (Han et al., 2021).

### 2.2 Heme reconstitution

A control heme reconstitution experiment was conducted as described previously (Chen et al., 2015). In brief, purified apo-*Dd*DyP from LB expression was mixed with hemin chloride with ∼1:2 molar ratio in a buffer containing 50 mM Tris-HCl, pH 7.0 and 150 mM NaCl, followed by incubation on ice for 30 min. The heme reconstituted holo-*Dd*DyP protein was then passed through a PD-10 desalting column (Cytiva, Sweden) to remove the excess hemin chloride.

### 2.3 UV-visible spectrophotometry

All spectra were recorded on a Shimadzu UV-2700 PC spectrophotometer (Shimadzu Co., Japan) using a cuvette with 1.0 cm pathlength in a range of 200-700 nm at room temperature (20 ± 2.0 °C). For measurements purified *Dd*DyP was diluted with lysis buffer to a concentration of 0.4 mg ml^−1^ and the lysis buffer was used as a reference. All spectra were processed using the Origin software 2022b (OriginLab Co., USA).

### 2.4 Crystallization screening and crystal optimization of *Dd*DyP

Crystallization screening was performed using a NT8 Formulatrix robot (Formulatrix, USA). Hit was obtained from the C12 condition (20% PEG 6000, 0.1 M HEPES, pH 7.0, 0.01 M ZnCl_2_) of the PACT++ crystallization screen (Jena Bioscience, Germany) with 10 mg ml^−1^ of purified *Dd*DyP. This condition was further optimized to 15% PEG 6000, 0.1 M HEPES-NaOH, pH 7.0, and 0.01 M ZnCl_2_ crystallized with 15 mg ml^−1^ final concentration of purified *Dd*DyP, which gave rise to a maximum crystal size of 200 × 100 × 25 μm at 20 °C in 4-6 weeks. Crystals were harvested directly from the drops using nylon loops and flash-cooled in liquid nitrogen.

### 2.5 X-ray diffraction data collection, processing and structure determination

X-ray diffraction datasets were collected at the P11/PETRA III beamline at DESY (Hamburg, Germany) using a flat focus with 20 × 20 μm^2^ (v × h) beam area at the sample position, 12.0 keV photon energy, and a photon flux of ∼2 × 10^10^ photon sec^−1^ and an exposure time of 100 ms for a total wedge of 360° with 0.1° oscillation recording step on EIGER 16M detector (Burkhardt et al., 2016). Data collection were performed at cryogenic temperature, 100K. Diffraction datasets were processed using the program XDS, and scaled with XSCALE in the XDS graphic user interface (Kabsch, 2010). The initial phase was obtained by molecular replacement using the *Dd*DyP peroxidase active structure (PDB ID: 7ODZ) as a reference model with the program Phaser in the phenix software (Afonine et al., 2012). The model was then refined in phenix and manually corrected in coot (Emsley et al., 2010). Radiation dose was estimated using the program RADDOSE-3D as described previously using the abovementioned parameters (Bury et al., 2018). For channel and cavity calculations *MOLEontile* tool (https://mole.uplo.cz/method) was used with the coordinate obtained from the final cycle of refinement (PDB ID: 8OHY) as a template (Sehnal et al., 2013). The interfaces of heme *b* and the oligomeric states analysis of *Dd*DyP were calculated using PISA (Protein Interfaces, Surfaces, and Assemblies) server (Krissinel and Henrick, 2007).

## 3 Results and Discussion

### 3.1 Heterologous expression and characterization of *Dd*DyP

For heterologous expression, a *Dd*DyP peroxidase gene was cloned into a pGEX-6P1 vector which has an HVR 3C excision site and a GST-tag in its N-terminal region as reported previously (Rai et al., 2021), however, we used the OverExpress *E. coli* C43(DE3) strain instead of BL21 for overexpression screening. *Dd*DyP was expressed at high and low temperature (37 and 20 °C) in LB and 2×YT medium with different yields, ranging from 2.1 and 14.1 mg of protein in average per 25 g of cells, respectively (Table 1). Figure 1A-B shows the SDS-PAGE analyses of the typical expression and purification of *Dd*DyP in *E. coli* C43(DE3) strain. Protein expressed at 37 °C, however, has transparent to pale brownish colour, whereas those of 20 °C exhibited a darker brownish colour (Figure 1C). The proteins purified from high and low temperature have a reasonable purity with less aggregation (Figure 1) with ∼35 kDa molecular weight as confirmed by the SDS-PAGE analysis. However, no crystallization hit was obtained from these conditions despite several attempts with various crystallization screens (Table 1). Since that our purified *Dd*DyP has sufficient purity for crystallization, yet we did not successfully crystallize it, we concluded that the instability of the protein during expression at 37 °C may be the cause for the unsuccessful crystallization. This is likely due to the improper protein folding, and thus lowering the temperature of the expression may be one method for achieving a stable and correctly folded protein (Francis and Page, 2010; Huang et al., 2021). Indeed, when we expressed *Dd*DyP at 20 °C, it gave rise to a darker brownish protein (Figure 1C), a typical heme peroxidase colour of a native protein, especially when expressed in a 2×YT enriched medium, which is richer than LB. Intriguingly, *Dd*DyP expressed in 2×YT at lower temperature was the only condition that resulted in a successful crystallization yielding dark brownish crystals, an indication that heme *b* is preserved during purification and crystallization (Figure 1D). The 2×YT also yielded several times higher amount of protein than that obtained in LB (Table 1).

**Table 1.** Characterization of the native heme incorporation into *Dd*DyP expressed in *E. coli* C43(DE3) at different conditions, the protein yield and crystallization trials

| Overexpression | Soret peak (nm) | $R_Z$ value ($A_{\text{soret}}/A_{280}$) | Heme content (%)<sup>†</sup> | Protein yield (mg)<sup>‡</sup> | Crystallization hits |
| --- | --- | --- | --- | --- | --- |
| LB at 37 °C | 406 | 0.27 | 26.5 | 1.2 | No |
| LB at 20 °C | 405 | 0.502 | 49.3 | 2.96 | No |
| 2×YT at 20 °C | 402 | 0.93 | 90 | 14.1 | Yes |
| Reference* | 406 | 1.02 | 100 | 1.2 | Not tested |

\*The reference is the *DdDyP* with a reconstituted heme; from LB expression. <sup>†</sup>Heme content relevant to the reference value in this study, which was set to 100%. <sup>‡</sup>The yield is normalized to 25.0 g of wet weight of overexpressed cells used for purification.

**Figure 1.**
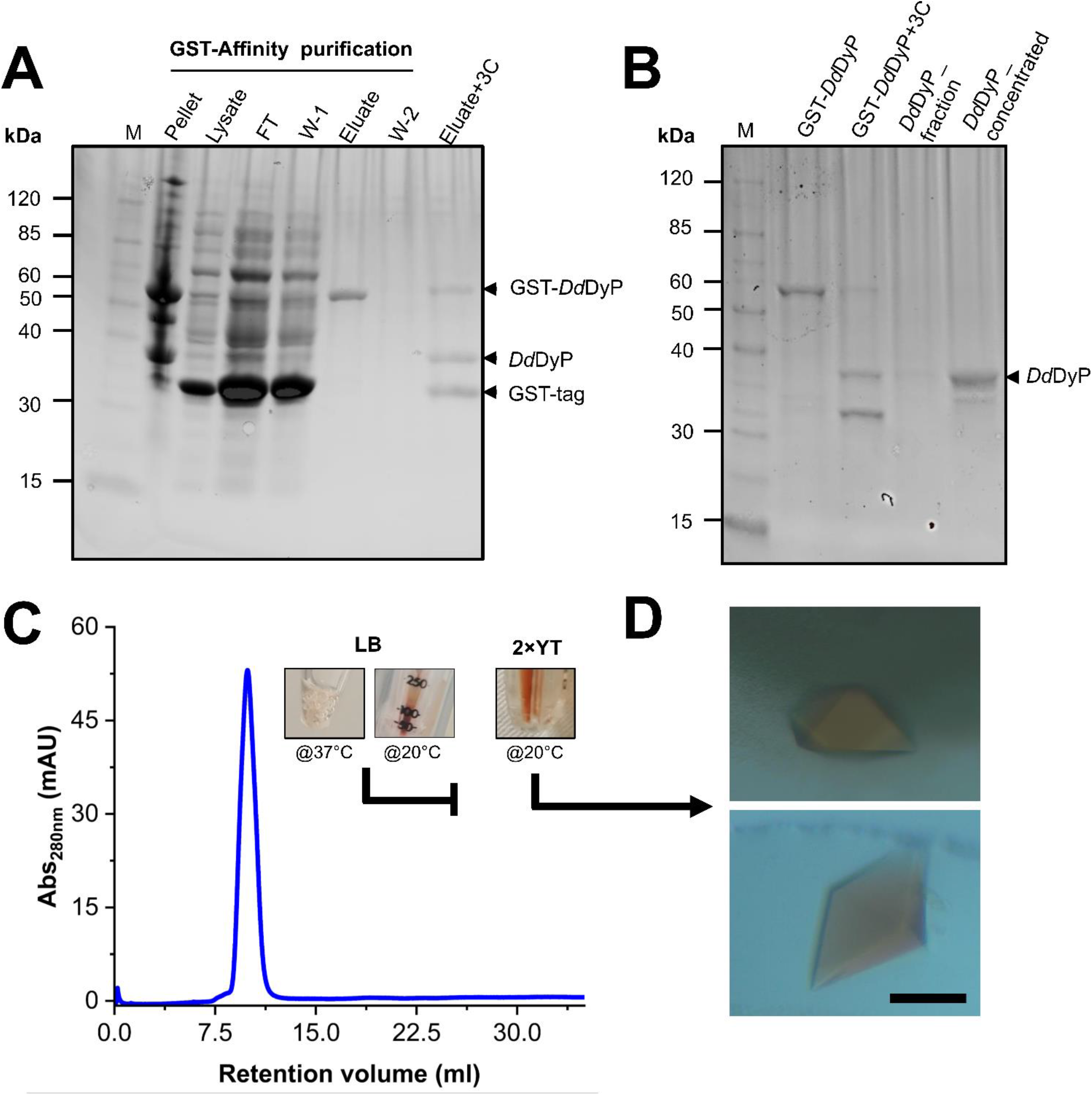
Purification, characterization and crystallization of *Dd*DyP. (A) An SDS-PAGE showing the glutathione sepharose affinity purification. Lane M, protein marker; FT lane, flow-through of the unbound proteins from the lysate; W-1 lane, step 1 washing of the Glutathione Sepharose column; and W-2, the second washing step of the column after elution (B) the HRV 3C digestion of GST-*Dd*DyP expressed protein complex and GST-free *Dd*DyP protein. (C) The Gel-filtration profile of purified *Dd*DyPs and representative samples from 2×YT expressed at 37 and 20 °C, and LB at 37 °C, and (D) brownish crystals from two different purification batches of independently overexpressed *Dd*DyP protein. The scale bar in “D” represents 100 μm.

To examine the quality of the electronic absorbance of purified *Dd*DyP we used a UV-visible spectrophotometer. Figure 2 shows the spectral analysis of the purified *Dd*DyP protein from different conditions. *Dd*DyP shows weak absorbance Soret peak at λ = 405 nm and a Reinheitszahl (*R*_Z_) (*A*_soret_/*A*_280_) value of ∼0.27 when expressed in LB at 37 °C (Table 1). This *R*_Z_ value is about 2 times higher than that obtained previously on peroxidases that were expressed using BL21(DE3) strain (Rai et al., 2014; Fiege et al., 2018). As shown in Table 1, we observed that the *R*_Z_ value increases ×2–3 times to reach ∼0.93 with λ = 402 nm of the Soret peak when expressing *Dd*DyP in enriched 2×YT medium at low temperature (20 °C). This value is comparable to our heme reconstitution reference (Figure 2E) and significantly higher (about 7 times) than those previously reported, when DyP-peroxidases were expressed without adding heme supplements or *δ*-ALA during expression (Fiege et al., 2018; Krainer et al., 2015). Heme biosynthesis in *E. coli* relies on an L-glutamate amino acid and other proteinoid biochemicals which are abundant in both tryptone and yeast extracts—major components in 2×YT and LB medium (Layer et al., 2010; Krainer et al., 2015). The 2×YT medium has a double amount of tryptone and yeast extracts comparing to LB medium. This might explain the higher level of heme biosynthesis, and thus the native incorporation into *Dd*DyP, in 2×YT than that in LB (Table 1). The heme reconstituted *Dd*DyP from LB expression shows a Soret absorbance at 406 nm and an electron transfer (ET) (*Q*-band) at λ = 497 nm plus two additional ET bands at 536 nm and 576 nm as well as a charge transfer (CT) component at 636 nm, preserving some bacterial peroxidase features (Chen and Li, 2016). This region differs significantly from that previously reported in *Dd*DyP, which showed an ET and CT band at 506 nm and 636 nm, respectively (Rai et al., 2021). We also observed that the *Dd*DyP with a natively incorporated heme has a broad ET peak with *λ*_max_ = 508 nm, which is slightly red shifted with Δ*λ* = 9 nm and Δ*λ* = 2 nm, comparing to that of the ET bands of the reference (Figure 2) and a previous work, respectively (Rai et al., 2021). The *Q*-band region also reveals a unique shoulder at the ET band with 567 nm absorbance (Figure 2C).

**Figure 2.**
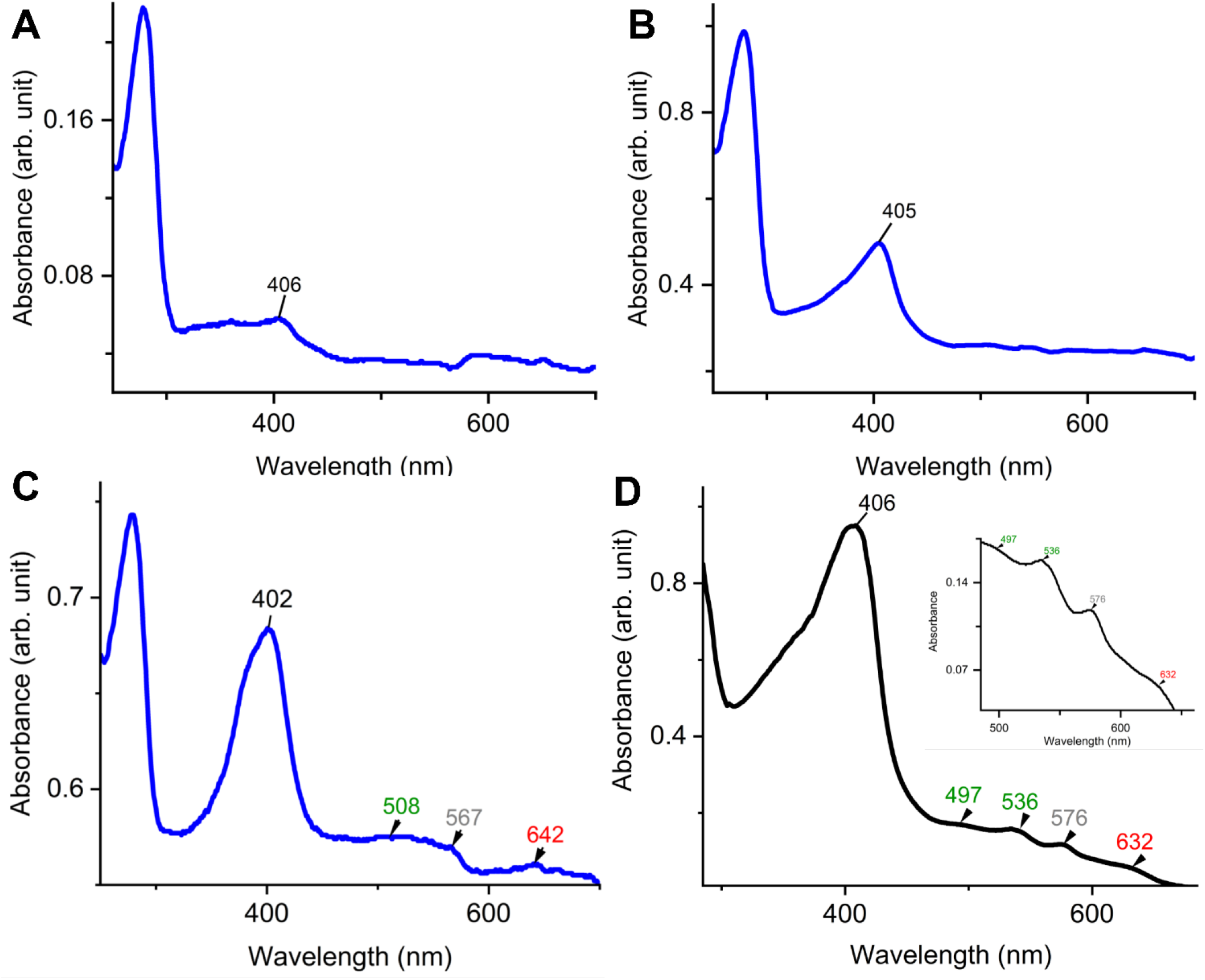
UV-visible electronic absorbance of *Dd*DyP of different overexpression conditions. (A) a typical spectrum of a purified *Dd*DyP when expressed in LB medium at 37 °C, (B) and (C) *Dy*DyP electronic spectra when expressed at 20 °C in LB and 2×YT medium, respectively. The ET *Q*-band and charge transfer band in the region from 500–700 nm of the 2×YT condition are described in (C). (D) The reference UV-visible spectra of LB *Dd*DyP after reconstitution with hemin chloride, and the small box in (D) represents a magnified view of the *Q*-band and charge transfer regions from 500–685 nm.

### 3.2 Native structure of *Dd*DyP peroxidase bound to a water molecule

Several *Dd*DyP structures have been resolved so far including a single native structure that is in complex with cyanide (PDB: 7O9L) (Rai et al., 2021), however there is no available structure that describes the native resting state. Here, to get insight into the heme binding pocket in its native form, we crystallized the native *Dd*DyP peroxidase bound to a natively incorporated heme and compared it with that resolved in complex with cyanide. A single crystal with a size of 150 × 80 × 30 μm^3^ size was used for diffraction data collection (Figure 1D). The crystal data collection and refinement statistics are shown in Table 2. Native *Dd*DyP peroxidase is crystallized in a tetragonal space group *P*4_1_ 2_1_ 2 similar to previously reported structures (Rai et al., 2014), with exception that the X-ray data of the current crystal condition can be equally processed and resolved in an additional space group (*P*4_3_ 2_1_ 2) (Table 2). Moreover, the crystal unit cell exhibited significantly shorter axes, giving rise to about 35% smaller cell volume with 52.8% solvent content and 2.62 Å^3^ Da^−1^ of Matthew’s coefficient (*V*_m_), indicating the presence of a single molecule per asymmetric unit. The solvent content is decreased by ∼20% comparing to that previously reported (Rai et al., 2021). This is more likely due to a relatively high concentration of the dehydrating precipitant (∼30% PEG 6000), as previously reported in other systems (Umena et al., 2011; Koua et al., 2013). Such high PEG concentration causes a shrinking in the protein crystals by dehydration which shorten the axes and leads to a tightly packed unit cell (Figure S1). The average radiation dose on a single crystal was estimated with RADDOSE-3D (Bury et al., 2018) to be ∼0.58 MGy (Gy = J•kg^-1^) (Table 2) which lies well below the 20 MGy dose limit suggested by Henderson (Henderson, 1990) or the 30 MGy suggested by Owen and Garman (Owen et al., 2006), indicating that the structure is less affected by radiation damage.

**Table 2.** X-ray diffraction data collection and crystallography refinement statistics

|  | Native *DdDyP* structure |
| --- | --- |
| PDB ID | 8OHY |
| **Data Collection** |  |
| Beamline | P11/PETRA III at DESY |
| Photon Energy (keV) | 12.0 |
| Photon flux (ph . s<sup>-1</sup>) | ~2 × 10<sup>10</sup> |
| Radiation dose (MGy) | ~0.58 |
| Space group | *P*4<sub>1</sub> 2<sub>1</sub> 2 |
| Cell dimensions |  |
| a, b, c (Å) | 99.88 99.88 73.12 |
| α, β, γ (°) | 90.00 90.00 90.00 |
| Resolution (Å) | 44.67–1.95 (2.02–1.95)* |
| *R*<sub>merge</sub><sup>†</sup> | 0.2715 (4.643)* |
| *I*/σ*I* | 12.02 (0.6)* |
| Completeness (%) | 98.54 (91.55)* |
| Multiplicity | 26.7 (26.0)* |
| *CC*<sub>1/2</sub> | 0.999 (0.411)* |
| *CC* * | 1.0 (0.764)* |
| Wilson *B*-factor (Å<sup>2</sup>) | 36.8 |
| **Refinement** |  |
| Resolution range (Å) | 44.67–1.95 |
| No. of reflections (unique) | 27129 (2471)* |
| *R*<sub>work</sub> / *R*<sub>free</sub><sup>‡</sup> | 0.206 (0.400)* / 0.247 (0.445)* |
| No. of atoms | 2662 |
| Protein | 2459 |
| Ligands | 74 |
| Solvent | 163 |
| No. of residues | 306 |
| Average *B*-factor (Å<sup>2</sup>) | 41.29 |
| Protein | 41.23 |
| Ligands | 39.63 |
| Solvents | 42.76 |
| r.m.s. deviations |  |
| Bonds (Å) | 0.005 |
| Angles (°) | 0.71 |
| Ramachandran (%) |  |
| Favored | 98.03 |
| Allowed | 1.97 |
| Outliers | 0.00 |
| Rotamer outliers (%) | 0.00 |
| Clashscore | 1.00 |

\*Values in parenthesis are of the highest resolution shell. <sup>†</sup>*R*<sub>merge</sub> = $\sum_h \sum_i |I_i(h) - \langle I(h) \rangle| /$ $\sum_h \sum_i |I_i(h)|$ , where *I<sub>i</sub>(h)* is the intensity measurement for a reflection *h* and $\langle I(h) \rangle$ is the mean intensity for this reflection. <sup>‡</sup>*R*<sub>work</sub> = $\sum_h ||F_{obs}| - |F_{calc}|| / \sum_h |F_{obs}|$ and *R*<sub>free</sub> was calculated using a randomly (5.0%) selected reflections.

The overall architecture of *Dd*DyP is similar to that of the typical DyP-type peroxidase superfamily (Sugano et al., 2007; Chen et al., 2015; Rai et al., 2021). *Dd*DyP contains a duplicated ferredoxin-like fold domain arranged as a *β*-barrel at the N- and C-terminals of the protein (Figure 3). It contains 12 *β*-sheets and 13 *α*-helices formed by 185 residues of the full chain (306 residues), and the remaining 121 residues involved in the formation of loop structures that link these secondary structures. Similar to all other known DyP-type peroxidases, *Dd*DyP contains *α*-helices with a unique β-sheet structure at the distal region of the heme plane (Strittmatter et al., 2013; Sugano et al., 2007). We determined the root mean square deviation (r.m.s.d.) between the C*α* (1-306 residues) of the present structure with that resolved in complex with cyanide (PDB ID: 7O9L) to be 0.18 Å, indicating the striking similarity between the two structures. Our PISA analysis predicted a stable dimer of *Dd*DyP in solution with 33 residues contributing to the dimer interface, similar to previously reported *Dd*DyP structures (Rai et al., 2021). These interfacial residues are distributed along the dimer interface from the N- to the C-terminal region. The dimeric structure reveals a solvent accessible area of 24290 Å^2^ and buried surface areas (BSA) of 5330 Å^2^, corresponding to about 22% of the total surface area of the protein. On the other hand, the BSA of the monomeric structure is 1341 Å^2^, corresponding to 9.6% of the total surface area of monomeric *Dd*DyP. It should be noted that our PISA analysis favoured a tetramer oligomeric state for the native cyanide *Dd*DyP (PDB ID: 7O9L) structure, displaying higher binding energy than that of the dimeric state. This indicates that *Dd*DyP protein may exist physiologically in various oligomeric states. Indeed, several DyP-type peroxidases have been reported to exist in different functional oligomeric states ranging from monomeric to tetrameric state (Yoshida et al., 2016; Pfanzagl et al., 2020; Liu et al., 2011; Zubieta et al., 2007). The catalytic arginine residue, Arg239 in *Dd*DyP, has been suggested to play a role in the protein oligomerization owing to its location and hydrogen bonding network with surface residues (Chen et al., 2015; Singh et al., 2012). In *Dd*DyP, Arg239 is buried in the hydrophobic cavity of the heme binding pocket, excluding its contribution in *Dd*DyP oligomerization. Moreover, our molecular replacement attempts aiming for a dimeric solution was not successful, thus we can reasonably conclude that our purified *Dd*DyP favours a monomeric state in crystal. It has been previously reported, based on sedimentation velocity analysis with analytical ultracentrifugation, that dimeric *Dd*DyP predominates in solution, which yielded a dimeric crystal structure (Rai et al., 2021). The crystal packing behaviour of the present structure (PDB ID: 8OHY) is significantly different from that described previously (Rai et al., 2021), likely due to a significantly low unit cell volume which exerts tight interactions between molecules in the unit cell (Figure S1). Note that the molecular contact within the unit cell is contributed by similar regions in both forms that is primarily the loop and β-sheet of the ferredoxin-fold like domain II at the C-terminal region (Figure S1).

**Figure 3.**
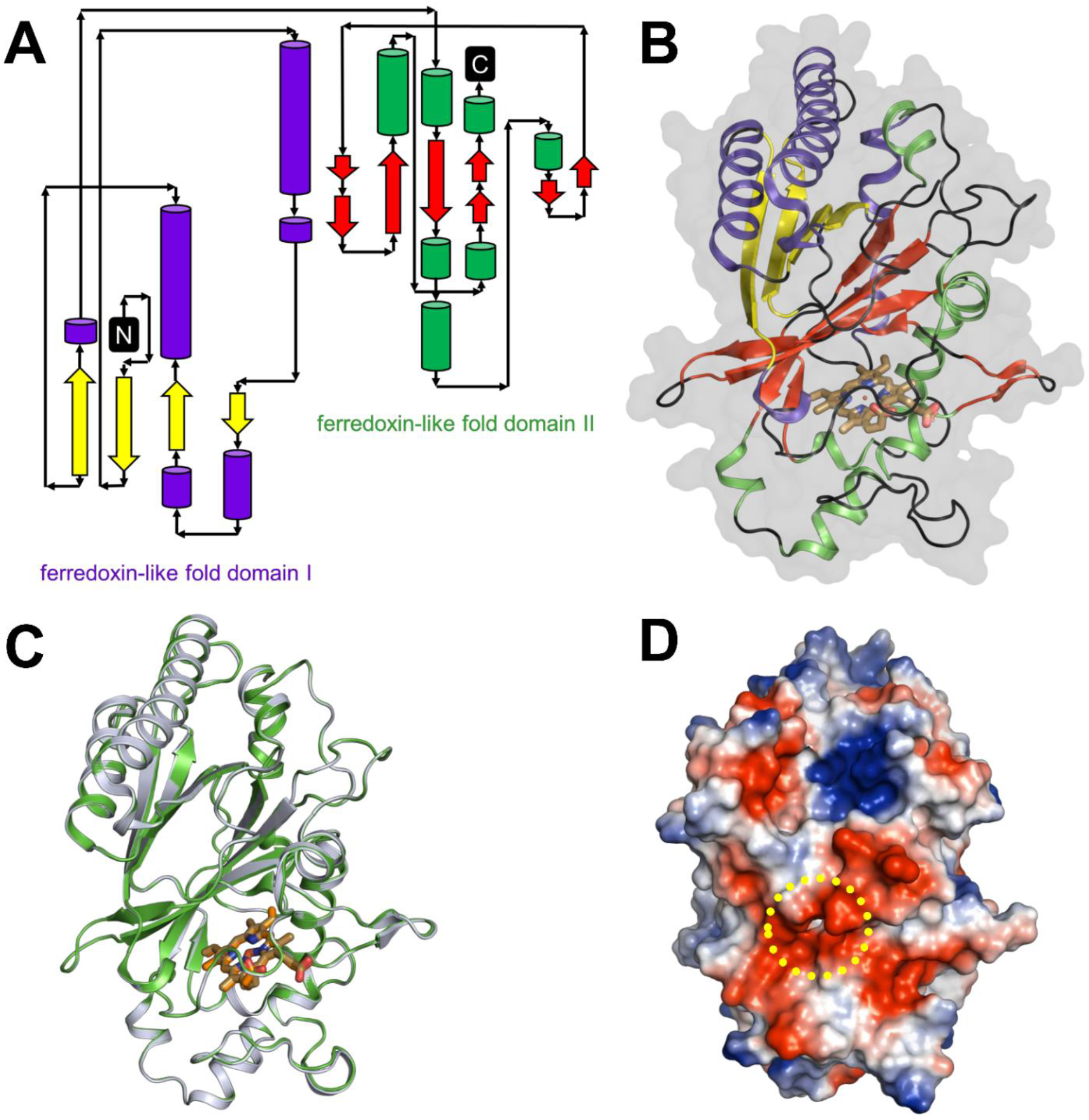
Overall structure of native *Dd*DyP and its superposition with native cyanide *Dd*DyP structure (PDB code: 7O9L). (A) Topology representation of the *Dd*DyP-type peroxidase. (B) Overall structure of monomeric *Dd*DyP showing the ferredoxin-like folds at the N- and C-terminals colored according to (A). (C) Superimposition of native *Dd*DyP structure (green) into a cyanide native structure (gray). (D) Electrostatic potentials surface of native *Dd*DyP colored from -5.7 kT (red) to +5.7 kT (blue) calculated using the program PyMOL (http://www.pymol.org/pymol). The yellow dashed circle highlights the heme binding pocket and possible pathway for H_2_O_2_ and/or substrate entry.

### 3.3 Geometry of a natively incorporated heme, its binding pocket and the implication in catalysis

Heme *b* in the DyP-type peroxidases, a protoheme IX, is either penta- or hexacoordinated (Sugano et al., 2007; Singh et al., 2012; Rodrigues et al., 2021). The native structure of *Dd*DyP accommodates heme *b* in a hydrophobic binding pocket flanked by the unique β-sheet at the distal side of the heme plane, the α-helices of the ferredoxin-like fold domain II (Figure 4) and a distinct long loop at its proximal side similar to previously reported DyP structures (Sugano et al., 2007; Zubieta et al., 2007; Liu et al., 2011). Our structural analysis shows that the native *Dd*DyP heme is hexacoordinated, of which the pyrrole rings of porphyrin contributed to tetradentate chelation via their nitrogen atoms and via the conserved His222 at the proximal side with a distance of 2.11 Å. The sixth coordination is provided by a water molecule (wat-184) with a distance of 2.79 Å, which is shorter by ∼0.1 Å than that of the reported Fe(III)-CN distance, indicating a stronger coordination to Fe(III) (Figure 4C). This distance is typical for Fe(III) of the resting state, implying that the model is unaltered by radiation damage (Chen et al., 2015). Wat-184 forms a strong hydrogen bond (∼2.2 Å) with wat-182 and a weaker hydrogen bond with the catalytic residue Arg239 (Figure 4C). Intriguingly, the environment of our native *Dd*DyP heme binding pocket is similar to that of the bacterial DypB peroxidases (Chen et al., 2015). We observed that wat-184 has slightly higher *B-*factor than wat-182, which may indicate its mobility and higher reactivity. On the other hand, wat-182 interacts via strong hydrogen bonds with the second catalytic residue Asp149 as well as Ser241, suggesting that these residues may act as proton acceptors to the H_2_O_2_ during the formation of compound I oxyferryl thereby contributing to its stabilization along with Arg239 (Figure 4C) as revealed in other A-type DyP peroxidases (Pfanzagl et al., 2018). The binding pocket is extensively occupied with water molecules which are in hydrogen bonding interaction with nearby residues that contribute to the heme stability (Figure 4C). Note that two of the water molecules (wat-150 and wat-159), which are in hydrogen bonding interaction with Asp149 and Arg137 near the heme access channel, substituted the 1,2-ethanediol molecule in the cyanide native structure that shifted Asp149 carboxylate group towards the cyanide, giving rise to increased *B*-factor of Asp149 comparing to its surrounding (Rai et al., 2021). This may indicate that this position is natively occupied by water molecules as demonstrated by our native structure.

**Figure 4.**
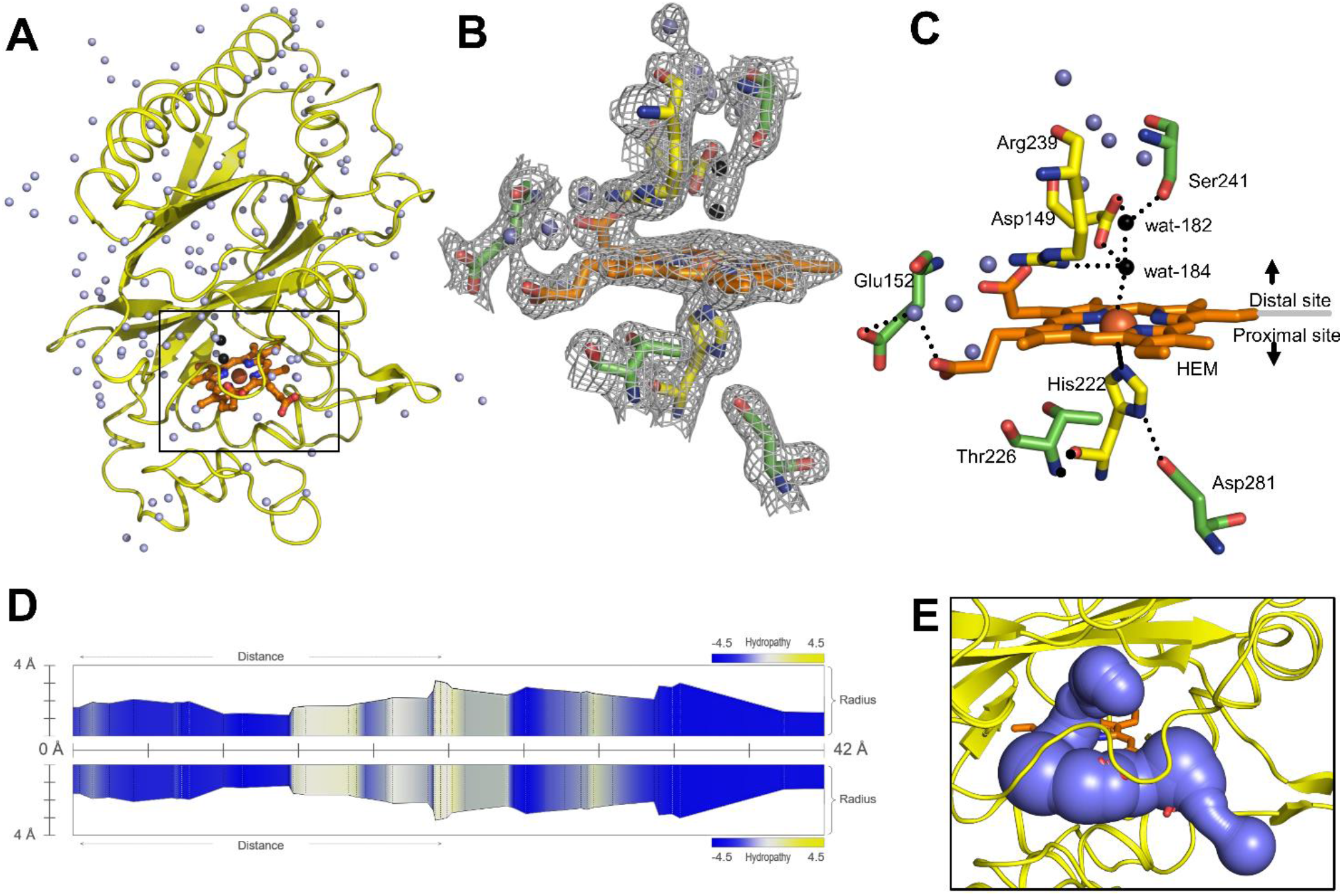
Monomeric structure of the native *Dd*DyP highlighting the heme binding pocket at a resolution of 1.95 Å (PDB code: 8OHY) and the channel characteristics of the system. (A) The overall monomeric structure of *Dd*DyP highlighting its heme binding pocket (black box). (B) The 2*m*Fo–*D*Fc electron density map (gray mesh) contoured at 1.0 sigma level displaying a magnified view of the heme binding pocket (active site). (C) The hydrogen bonding network (black dots) of the heme binding pocket at the distal and proximal sites of the heme plane. The two water molecules (wat-182 and wat-184) liganded to the heme-Fe(III) via hydrogen bond are highlighted in black, while the conserved key residues are displayed as yellow stick. All figures were generated using PyMOL software (http://www.pymol.org/pymol). (D) Two-dimensional representation of a large pore spanning the C-terminal region across the distal site of the heme binding pocket as a result of two channel convergence colored according to the hydropathy scores of the lining residues. (E) A 3D representation of the channel shown on (D).

The heme ligand is well resolved at a resolution of 1.95 Å as revealed by its 2*m*Fo–*D*Fc electron density map (Figure 4B and Figure S2), indicating unambiguous incorporation and binding of the heme in the apo-protein. This is an important finding as crystallization with purified *Dd*DyP proteins that have lower *R*_Z_ values were not successful, which might indicate that the heme on these purified proteins is not well accommodated in the binding pocket, affecting possibly their stability and hence the crystallization (see Table 1). The heme occupies 798 Å^2^ surface area, corresponding to 5.7% of the total surface area of the native structure, similar to that of the cyanide native *Dd*DyP structure and other *Dd*DyP structures (Rai et al., 2021). PISA analysis indicates that the solvation free energy gain (Δ^i^G) of the natively incorporated heme is -22.5 kcal mol^-1^ with 622 Å^2^ interface area comparing to an average of -22.3 kcal mol^-1^ for the reconstituted heme of the cyanide native structure (PDB ID: 7O9L) which has an interface interaction area of 609 Å^2^, indicating similar heme binding affinity with slightly better properties for the natively incorporated heme. The van der Waals interactions as well as the hydrogen-bonding network provided by nearby residues and water molecules may contribute to the stabilization of heme binding (Figure 4C) (Sacquin-Mora and Lavery, 2006; Mogharrab et al., 2007). Our analysis shows that the heme is stabilized, along the plane, via its carboxylate oxygens by hydrogen bonding with several water molecules (wat-120, wat-79 and wat-166), and three residues, Glu152, Arg204 and Arg239. The Arg239 interacts weakly with the heme carboxylates oxygens via two hydrogen bonds with a distance of 3.3-3.4 Å, whereas Glu152 and Arg204 form hydrogen bonding with the heme via wat-120 and wat-166, respectively. These interactions indicate that the heme is well stabilized in our model (PDB ID: 8OHY), which confirms the correct geometry of its native incorporation, yielding comparable binding pocket geometry to that prepared with *in vitro* reconstitution (Chen et al., 2015; Rai et al., 2021). Furthermore, the superimposition with the cyanide native structure indicates a striking similarity (r.m.s.d. = ∼0.18 Å) around the heme binding pocket including the flanking loop at the proximal side of the heme plane (residues 204-220) (Figure 3D). A side-specific mutagenesis study in DypB found that this proximal loop may have significant role in the heme stability (Rodrigues et al., 2021). This loop has also been implicated in the stabilization of the substrate owing to its flexibility thereby facilitating the substrate/product turnover by flipping in and out around the heme binding site (Liu et al., 2011).

Further, we used the *MOLEonline* tool (https://mole.upol.cz/online) to analyse the cavities and tunnels nearby the heme binding pocket and those in long-range distances (Figure S3) (Sehnal et al., 2013). Overall, 14 tunnels were identified, three of which are located next to the heme and perpendicular to each other with characteristics that might have an implication in the *Dd*DyP catalysis—entry of substrates and exit of reaction products. These channels may serve as entry gates for H_2_O_2_ thereby facilitating the enzyme activation required for the oxidative catalysis (Chen et al., 2015; Yoshida et al., 2016; Habib et al., 2019). Two tunnels have average diameter of ∼3.0 Å, which is sufficient to facilitate the entry of H_2_O_2_ and perhaps the exit of reaction products of similar size. All tunnels are lined with hydrophobic residues in the middle of the channel as well as several key catalytic residues in the distal and proximal sides of the heme plane. In particular, Arg239, Asp149, His222, Glu152, Ser241, and Thr226 in addition to several hydrophobic residues where found in two proximal channels (Figure S3), which are converged to form a main channel with a length of 42 Å and a diameter of ∼4.0 Å. The access gate of this channel is lined with charged residues as shown in Figure 4D and E, indicating its implication in the substrate entry. Further, we identified two major cavities at the distal side of the heme plane, of which one cavity (cavity 1) has a volume of 2881 Å^3^, corresponding to 9.6% of the total surface of *Dd*DyP and 4.7 times of the heme molecule. It is located at the heme binding pocket, accommodating the main channel at the binding pocket and extends to the proximal side of the heme plane, indicating a role for this cavity in the catalysis of DyP-peroxidases (Rai et al., 2021; Habib et al., 2019; Yoshida et al., 2011). The second cavity (cavity 2) with approximately half a volume of that of cavity 1 (1411 Å^3^) is located at the N-terminal region distant from the heme binding pocket and in contact with cavity 2 near the molecular centre of *Dd*DyP (Figure S3). The presence of such cavities is important for accommodating wide-range of substrates thereby fulfilling the substrate broad specificity of DyP-type peroxidases (Pfanzagl et al., 2019; Silva et al., 2023).

## 4 Conclusions

In conclusion, we demonstrated the use of *E. coli* C43(DE3) strain for heterologous expression of *Dd*DyP peroxidase, without the use of a heme precursor *δ*-ALA, hemin chloride or iron supplement, to produce to produce *Dd*DyP holoprotein with a natively incorporated heme, relying primarily on the *E. coli* heme biosynthesis by benefiting from the use of enriched medium and low temperature during expression, which yielded an *R*_Z_ value of ∼1.0 and a holoprotein with sufficient stability. We further showed, by mean of X-ray crystallography, that the native *Dd*DyP expressed in this condition has comparable heme geometry and binding properties. Our study also demonstrates that the natively incorporated heme is well stabilized via hydrogen bonds provided by nearby Arg239, Glu152 and water molecules in addition to van der Waals interactions between the porphyrin rings and surrounding residues within van der Waals distances. Two cavities occupying a total volume of 4292 Å^3^, corresponding to 14.3% of the total monomeric volume (29951 Å^3^), were identified. Of which the main cavity (cavity 1) around the heme binding pocket was found to accommodate a large access channel that spans the heme binding pocket. Moreover, the high-quality crystals optimized in this work would be suitable for use as a model for metalloenzymes to study the dynamics and substrate binding kinetics during catalysis. This can be achieved by, for instance, the mixing-and-inject time-resolved serial femtosecond crystallography approach (Pandey et al., 2021), which enables tracking the formation of the reaction intermediates as well as the mechanism of substrate breakdown into products as demonstrated in other metalloproteins (Worrall and Hough, 2022; Malla and Schmidt, 2022).

## 6 Author Contributions

Conceptualization, F.H.M.K.; experiments, Ö.K., L.B., H.H., and F.H.M.K.; formal analysis and data interpretation, F.H.M.K.; Diffraction data collection and processing, F.H.M.K., Ö.K., and S.K.; X-ray structural analysis, F.H.M.K.; contributed funding/reagents/analytic tools, A.P.M; supervised the project, F.H.M.K.; manuscript writing, F.H.M.K. with input from all authors. All authors read and approved the final version of this manuscript.

## 8 Data availability statement

The crystal structure data was deposited to the protein data bank under the PDB ID: 8OHY and is publicly available under the PDB digital object identifier http://doi.org/10.2210/pdb8OHY/pdb.

## 9 Acknowledgments

We would like to thank the staff members of the SPB/SFX instrument of European XFEL GmbH for fruitful discussion during the conduction of this research. We are grateful to the staff members of the XBI Biolab at European XFEL GmbH for technical support. We acknowledge the use of the XBI Biolab at European XFEL GmbH, enabled by the XBI User Consortium. X-ray diffraction experiments at the P11/PETRA III beamline in DESY were carried out via the proposal no. BAG-20211047 acquired by Huijong Han and Kristina Lorenzen from European XFEL GmbH. We also thank the staff members of the P11/PETRA III beamline at DESY, Hamburg. We thank Richard Bean for the critical reading of the manuscript.

## SCHEME AND FIGURE LEGENDS

**Scheme 1.**
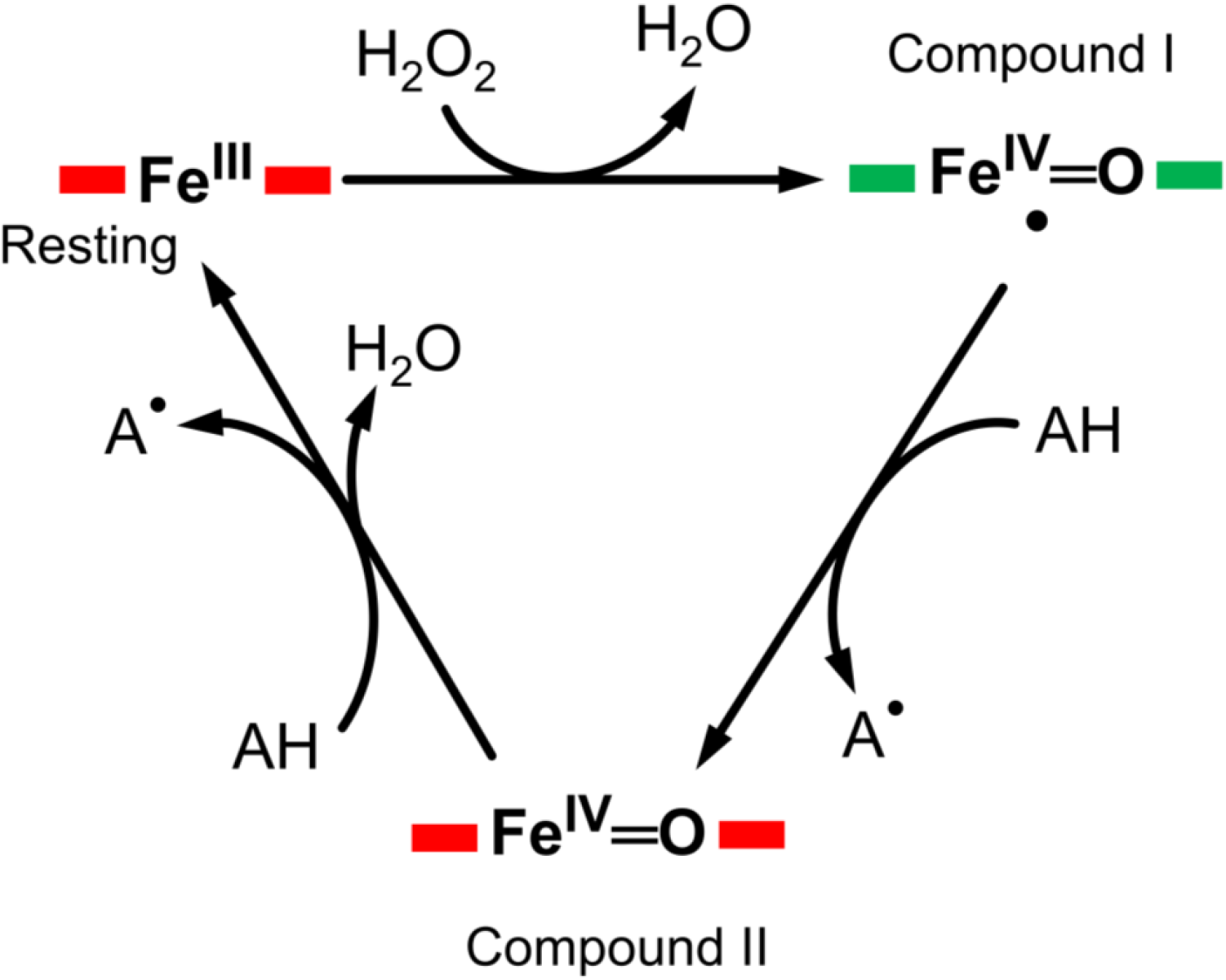
A typical enzymatic cycle of DyP-type peroxidases showing the interconversion between the resting state (ferric porphyrin), compound I heme oxoferryl species (porphyrin cationic radical) [Fe(IV)═O Por^•^]^+^; and compound II intermediate state, [Fe(IV)═O]^+^. The AH is the reducing substrate which is oxidized into an intermediate radical product (A^•^) during catalysis.

