## Supplemental Figure 1, Figure 2 and Figure 3 for "Heterologous expression, purification and structural features of native *Dictyostelium discoideum* dye-decolorizing peroxidase bound to a natively incorporated heme"

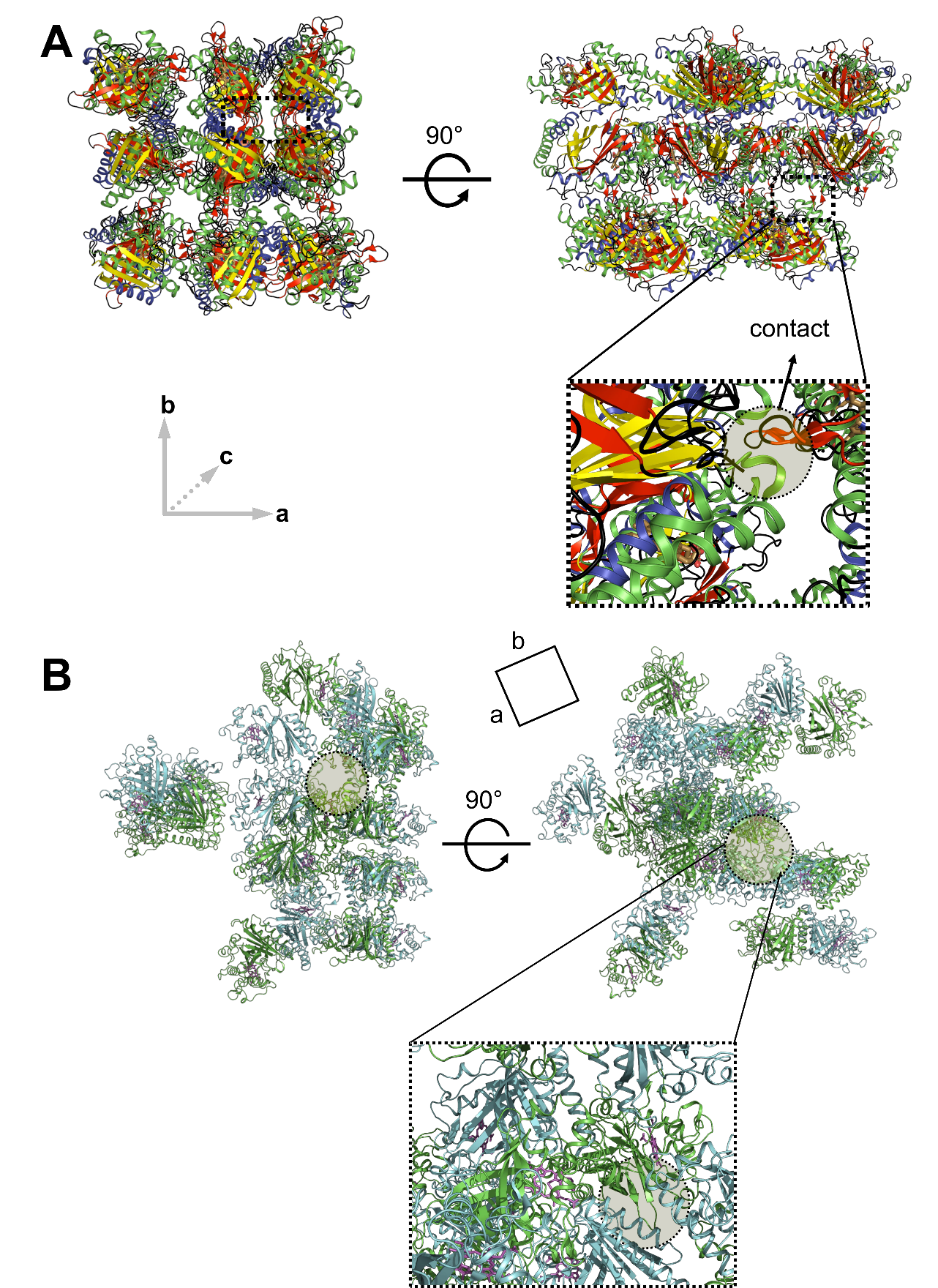

**Supplementary Figure 1.** Comparison of the *Dd*DyP crystal packing pattern of the monomeric structure of (A) this study (PDB ID: 8OHY) and (B) the dimeric structure (PDB ID: 7O9L). Crystal contact regions are highlighted with dashed rectangular (A) or dashed circle (B).

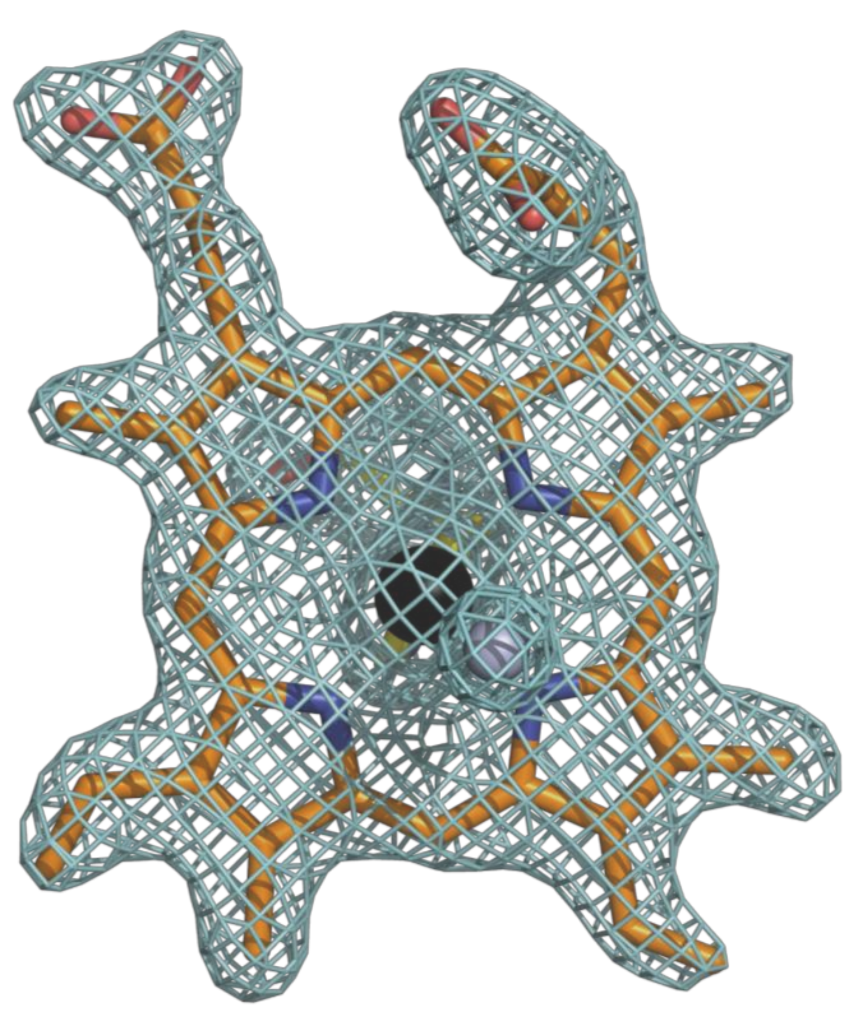

**Supplementary Figure 2.** Quality of heme *b* ligand viewed from the distal side showing the 2m*F*o–D*F*c countered at 1.25 sigma level.

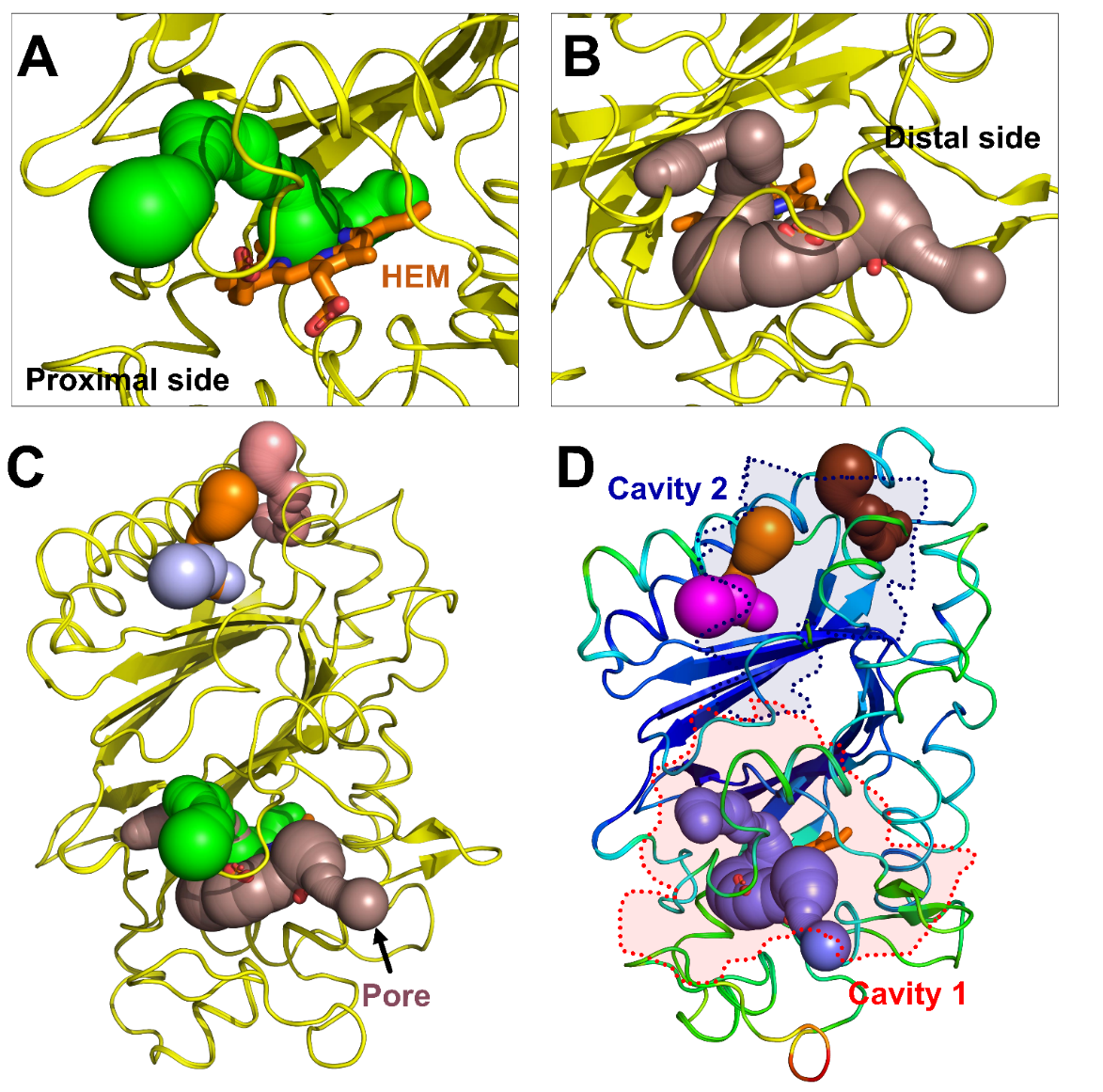

**Supplementary Figure S3.** Predicted tunnels and cavities in the native *Dd*DyP structure. (A) and (B) are the 3D representation of the three different tunnels which are in close proximity to the heme binding pocket. (C) Shows a main pore as resulted from the convergence of two channels at the heme binding pocket and the set of distant channels at the N-terminal site. (D) The locations of 2 predicted cavities highlighted with dashed line (red color) for cavity 1 enclosing the binding pocket including the main channel and dashed line with blue color for cavity 2 at the N-terminal site—a long-range cavity relative to the heme binding pocket.
